# Direct capture of CRISPR guides enables scalable, multiplexed, and multi-omic Perturb-seq

**DOI:** 10.1101/503367

**Authors:** Joseph M. Replogle, Albert Xu, Thomas M. Norman, Elliott J. Meer, Jessica M. Terry, Daniel Riordan, Niranjan Srinivas, Tarjei S. Mikkelsen, Jonathan S. Weissman, Britt Adamson

**Affiliations:** Tetrad Graduate Program, University of California, San Francisco, San Francisco, CA 94158, USA; Medical Scientist Training Program, University of California, San Francisco, San Francisco, CA 94158, USA; Department of Cellular and Molecular Pharmacology, University of California, San Francisco, San Francisco, CA 94158, USA; Howard Hughes Medical Institute, University of California, San Francisco, San Francisco, CA 94158, USA; California Institute for Quantitative Biomedical Research, University of California, San Francisco, San Francisco, CA 94158, USA; 10x Genomics Inc., Pleasanton, California, 94566, USA; Present address: Lewis-Sigler Institute for Integrative Genomics, Princeton University, Princeton, NJ 08544, USA; Present address: Department of Molecular Biology, Princeton University, Princeton, NJ 08544, USA

## Abstract

Pairing CRISPR-based genetic screens with single-cell transcriptional phenotypes (Perturb-seq) has advanced efforts to explore the function of mammalian genes and genetic networks. We present strategies for Perturb-seq that enable direct capture of CRISPR sgRNAs within 3’ or 5’ single-cell RNA-sequencing libraries using the 10x Genomics platform. This technology greatly expands the accessibility, scalability, and flexibility of Perturb-seq, specifically enabling use with programmed combinatorial perturbations and multiplexing with multi-omic measurements.

## Main

Two major advances have propelled efforts to define gene function in mammalian cells: CRISPR/Cas9-based genetic approaches that allow any gene or gene combination to be turned up, down, or off, and methods for massively parallel, high-resolution profiling of the phenotypic consequences of these changes. The pairing of pooled CRISPR screens with single-cell gene expression profiling, called Perturb-seq^1-5^, is particularly powerful as it enables principled and unbiased definition of gene function and systematic delineation of complex molecular networks^6^. Moreover, Perturb-seq presents an approach for tackling problems that were previously challenging or intractable, including identifying the mechanisms of action of small molecule drugs and natural products, synthetic lethal vulnerabilities in tumor subpopulations, genetic factors that induce cellular differentiation and reprogramming, and suppressors of inherited genetic diseases with complex cellular phenotypes. However, current implementations of Perturb-seq face technical limitations that make them difficult to pair with emerging single-cell technologies and hinder their use with programmed combinatorial perturbations. These limitations stem from a fundamental requirement of all previously published platforms: the use of indexing transcripts to assign perturbation identity to single-cell phenotypes.

To enable the monitoring of CRISPR guide RNAs (sgRNAs or guides) on standard platforms for droplet-based single-cell RNA-sequencing (scRNA-seq)^7-9^, which typically capture only the 3’ ends of polyadenylated RNAs (**Supplementary Fig. 1a**), current implementations of Perturb-seq use specialized expression vectors to co-express polyadenylated indexing transcripts alongside non-polyadenylated sgRNAs (**Fig. 1a**). For example, in an approach we developed together with the Regev lab (hereafter referred to as GBC Perturb-seq), indexing transcripts are programmed with unique guide barcodes (GBCs)^1,3^. When paired with specific amplification protocols that allow for deep sequencing, these highly expressed transcripts enable near perfect assignment of guide identities to individual cells. However, sgRNA-GBC uncoupling, which can occur during lentiviral transduction of co-packaged sgRNA-GBC libraries, corrupts guide assignment to cells^10-13^. GBC-based platforms are therefore best suited for use with libraries made by arrayed packaging, which limits scalability, unless specific steps are taken to mitigate crossovers^12,13^. An alternate implementation of Perturb-seq, termed CROP-seq, minimizes the problem of barcode uncoupling^4^. This approach relies on a clever expression vector to faithfully pair the expression of indexing transcripts encoding sgRNA sequences (cloned into the CROP-seq vector) with the expression of sgRNAs (copied by a duplication event during lentiviral transduction). Because this pairing is not confounded by recombination, CROP-seq is compatible with pooled library generation; however, CROP-seq has other constraints. Mainly, features of CROP-seq’s indexing strategy limit the method’s utility for exploring combinatorial perturbations between defined sets of gene pairs. Therefore, across implementation strategies, reliance on indexing transcripts (and the specialized vector systems they necessitate) limits the scale or application of Perturb-seq. Here, we overcome these limitations by establishing methods for scRNA-seq with independent capture of expressed guide RNAs, a streamlined technology, called “direct capture Perturb-seq”, that greatly expands the accessibility, scale, and utility of single-cell molecular screens.

**Figure 1:**
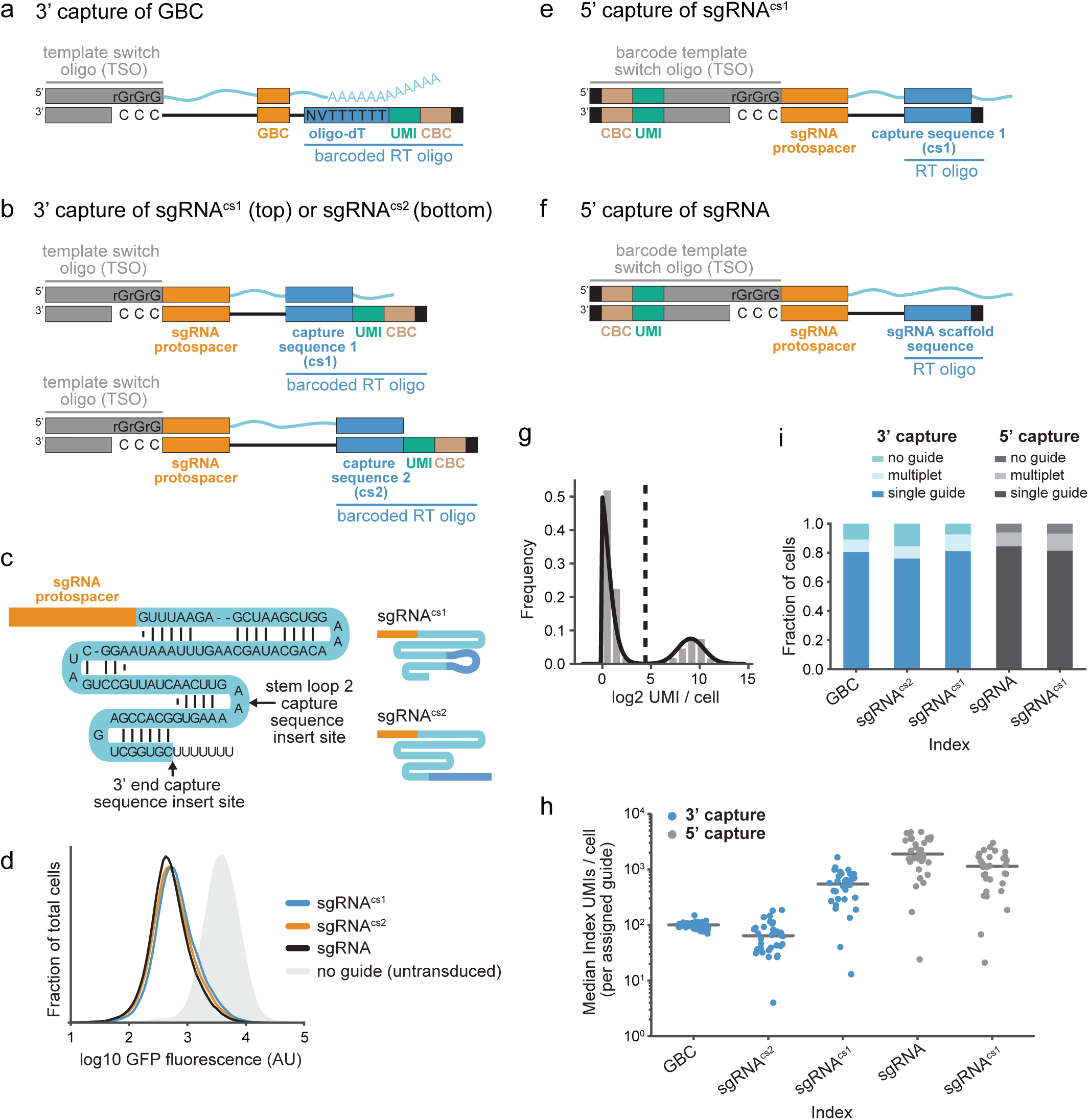
Direct capture of CRISPR guides on platforms for 3’ and 5’ single-cell RNA-sequencing. **(a)** Schematic of guide barcode (GBC) capture during Perturb-seq by 3’ scRNA-seq. GBC-containing mRNA (top) is reverse transcribed into indexed cDNA (bottom) using barcoded oligo-dT. UMI, unique molecular identifier. CBC, cell barcode. **(b)** Schematic of guide capture using integrated capture sequences (cs1 and cs2) during Perturb-seq by 3’ scRNA-seq. Capture sequences within the constant regions of two guide RNAs (top) anneal to barcoded, target-specific RT primers. Indexed DNA (bottom) is produced by reverse transcription. **(c)** Schematic of guide RNAs. Arrows indicate the positions of capture sequence insertions. Upper right: sgRNA^cs1^ with capture sequence 1. Lower right: sgRNA^cs2^ with capture sequence 2. **(d)** Gaussian kernel density estimates of normalized flow-cytometry measurements representing GFP expression demonstrate CRISPRi activity of the indicated guide RNAs (programmed with the GFP-targeting protospacer EGFP-NT2). **(e)** Schematic of guide capture using an integrated constant sequence during Perturb-seq by 5’ scRNA-seq. A capture sequence (cs1) within the constant region of sgRNA^cs1^ (top) anneals to an unbarcoded, target-specific RT oligo. Indexing of reverse transcribed DNA (bottom) occurs after template switch. **(f)** Schematic of guide capture without an integrated constant sequence during Perturb-seq by 5’ scRNA-seq. A guide RNA containing a standard constant region (top) anneals to a guide-specific RT oligo. Indexing of reverse transcribed DNA (bottom) occurs after a template switch. **(g)** Representative guide identity mapping corresponding to NegCtrl3 in Perturb-seq experiment conducted by 3’ sgRNA^cs1^ capture. Mapping relies on fitting a 2-component Poisson and Gaussian mixture model (black line), where cells with a posterior probability >0.5 (dotted line) of belonging to the upper mode component are assigned to NegCtrl3. **(h)** Index (GBC or guide) capture rates per cell across experiments conducted with GBC Perturb-seq and direct capture Perturb-seq. Data represent median index UMI counts per cell for cells bearing each of 32 protospacers across platforms. Grey lines indicate median values. **(i)** Index (GBC or guide) assignment rates across experiments conducted with GBC Perturb-seq and direct capture Perturb-seq. The fraction of cells assigned no guide, a single guide, or more than one guide are indicated.

Massively parallel scRNA-seq is performed by partitioning individual cells into droplets for molecular barcoding by reverse transcription (RT)^7-9^. This RT affixes unique molecular identifiers (UMI) and cell barcodes (CBC) to the ends of polyadenylated mRNA molecules, typically using barcoded oligo-dT to label 3’ ends (3’ scRNA-seq) (**Supplementary Fig. 1a**). Our first method for direct capture Perturb-seq implements 3’ scRNA-seq with parallel capture of non-polyadenylated guide RNAs (**Fig 1b** and **Supplementary Fig. 1b**). For this, we co-deliver barcoded, target-specific RT primers alongside similarly barcoded oligo-dT primers to droplet reactions. We deliver these barcoded oligos on gel beads (now commercially available from 10x Genomics in the Chromium Single Cell 3’ Reagent Kits v3 with Feature Barcoding technology), which we designed to carry two target-specific RT primers, each containing a single priming sequence or “capture sequence”. We optimized these sequences to satisfy three important criteria: (1) incorporation of the reverse complements of these sequences (also referred to as “capture sequences”) into sgRNAs (to allow for primer annealing during RT) should not compromise activity, (2) these sequences should facilitate robust and specific barcoding of targeted RNAs, and (3) addition of primers carrying these sequences to single-cell reactions should not interfere with the generation of gene expression profiles.

To identify capture sequences that satisfied the first of our criteria, we used CRISPR interference (CRISPRi)^14,15^ in K562s to screen candidate sequences for effects on guide activity conferred by their incorporation into one of two positions in the guide constant region: “stem loop 2” or the 3’ end (**Fig. 1c**). In agreement with previous observations^16^, we found that the extra sequence in the loop region minimally impacted guide activity, while incorporation of extra sequence near the 3’ end often decreased activity (**Supplementary Fig. 1c**). A notable exception was incorporation of polyA, which significantly compromised guide activity at either site. This generally discourages the use of polyadenylated guides with standard 3’ scRNA-seq for direct capture Perturb-seq. Our final capture sequences, capture sequence 1 (cs1) and capture sequence 2 (cs2), were chosen because cs1 incorporation into the loop, as contained in sgRNA^cs1^, and cs2 incorporation at the 3’ end, as contained in sgRNA^cs2^, do not compromise guide activity (**Fig. 1c,d** and **Supplementary Fig. 1c**). Notably, while the position of cs2 is interchangeable, incorporating cs1 at the 3’ end decreases guide activity (**Fig. 1d** and **Supplementary Fig. 1d**).

To maximize the generalizability of our method, we also developed direct capture Perturb-seq for 5’ scRNA-seq (**Fig. 1e,f** and **Supplementary Fig. 1b,e**). This complementary approach takes advantage of an existing droplet-based scRNA-seq platform (10x Genomics) that uses barcoded template switch oligos (TSO) and unbarcoded oligo-dT to affix indices to the 5’ ends of polyadenylated RNAs (**Supplementary Fig. 1e**). Unlike 3’ scRNA-seq, on this platform, oligo-dT RT primers are delivered to single-cell reactions as free oligos. Therefore, we were able to achieve capture of non-polyadenylated guide RNAs alongside global capture of polyadenylated mRNAs by mixing guide-specific RT primers with the standard RT enzyme mix containing oligo-dT. For this, we designed two guide-specific RT primers: one containing capture sequence 1 (cs1), designed to capture sgRNA^cs1^ (**Fig. 1e**), and another containing the reverse complement of a sequence from standard guide constant regions (**Fig. 1f**). We designed the latter primer to bind across many commonly-used guide variants making our approach amenable to screening across *Streptococcus pyogenes* Cas9 perturbation systems, including those with modified guide libraries such as SAM-based CRISPR activation (CRISPRa)^16^. In principle, our approach is also readily adaptable for use with guides from other species and for other classes of Cas proteins.

We next tested the performance of both approaches for direct capture Perturb-seq (3’ and 5’) and compared them to GBC Perturb-seq. For this, we screened three CRISPRi libraries, each targeting the same 30 genes with identical protospacer sequences (plus 2 non-targeting negative controls). These guides were chosen because depletion of the targeted genes had previously been shown to generate robust transcriptional signatures in K562s, in part due to activation of the unfolded protein response (UPR)^3^. To enable capture across Perturb-seq platforms, each library was built with a different guide constant region, specifically our parental constant region^17^, which was used in combination with our GBC Perturb-seq expression vector^3^, or the constant region from sgRNA^cs1^ (cs1 in the loop) or sgRNA^cs2^ (cs2 at the 3’ end) (**Fig 1c**). We designed and used specific amplification protocols for each experiment to deeply sequence our index molecules (GBC or guides) (**Supplementary Fig. 1b**). Promisingly, at a constant sequencing depth of 25 million aligned index reads per experiment, both of our direct capture platforms (3’ and 5’) demonstrated higher index capture (index UMIs per cell) than our GBC-based method (Mann Whitney U test: *p*=1e-9 for 3’ sgRNA^cs1^ capture; *p*=9e-10 for 5’ sgRNA^cs1^ capture; *p*=6e-11 for 5’ sgRNA capture) (**Supplementary Fig. 2a**), with the exception of 3’ capture of the sgRNA^cs2^ library, which had modestly lower capture (Mann Whitney U test: *p*=0.0002). Specifically, we observed median index UMIs/cell of 103.0 for GBC capture, 56.0 for 3’ sgRNA^cs2^ capture, 419.0 for 3’ sgRNA^cs1^ capture, 1593.5 for 5’ sgRNA capture, and 808.0 for 5’ sgRNA^cs1^ capture, demonstrating efficient capture of guide RNAs.

Next, we developed an approach to robustly assign guide identities to cells. For direct capture Perturb-seq, we did this by separating true guide-expressing cells from background cells by fitting a two-component mixture model to the log_2_-transformed guide UMIs per cell for each guide (**Fig. 1g**; see Methods). Background cells can arise (albeit at much lower levels) from spurious guide-CBC reads potentially generated from ambient guides during cell droplet formation or PCR chimeras. For both direct capture strategies (3’ and 5’), we found that guide capture rates vary considerably in a manner that was correlated across platforms (**Fig. 1h** and **Supplementary Fig. 2b,c**). Notably, these rates were related to the nucleotides at the 5’ ends of protospacers but not the protospacer GC content (**Supplementary Fig. 2d-f**). We observed that, for 90% of guides in each experiment, capture rates (guide UMIs/cell) fell within a range of 27-162 for 3’ capture of the sgRNA^cs2^ library, 92-1031 for 3’ capture of the sgRNA^cs1^ library, 349-4566 for 5’ capture of the standard sgRNA library, and 132-2357 for 5’ capture of the sgRNA^cs1^ library. However, the effect of this variability was mitigated by the overall high capture rates (**Fig. 1h**), and our assignment procedure was able to robustly assign guide identities to 84-94% of cells for our direct capture methods (compared to 89% for GBC Perturb-seq) with roughly expected guide distributions across all platforms (**Fig. 1i** and **Supplementary Fig. 2g**). These assignment rates tracked the true percentages of guide-bearing cells (97% for the standard sgRNA library, 94% for the sgRNA^cs2^ library, and 96% for the sgRNA^cs1^ library) as determined by BFP expression (which reports on vector integration). Importantly, we were also able to robustly identify cells with multiple guides at rates consistent (within 1-1.6 fold) with our expectations from library transduction (assumed Poisson infection distribution) and droplet cell capture (published multiple encapsulation rates) (**Fig. 1i**).

Given our ability to confidently assign guide identities to cells, we next sought to test the performance of direct capture Perturb-seq to study gene function. For this, we focused on the 3’ and 5’ platforms with the highest guide assignment rates: Perturb-seq by 3’ sgRNA^cs1^ capture and by 5’ capture of standard sgRNAs. High-content Perturb-seq phenotypes enable (1) high precision functional clustering of target genes, (2) identification of transcriptional phenotypes caused by individual perturbations, and (3) delineation of gene expression regulons. To address the first of these, we hierarchically clustered the perturbations in each of our experiments based on their pseudo-bulk expression profiles, which we generated by averaging normalized mRNA counts across all cells bearing the same guide RNA (**Fig. 2a**). This clustering recapitulated known functional and physical interactions between the targeted genes, including those among translation initiation factors (EIF2B2/EIF2B3/EIF2B4), UFMylation components (UFM1/UFL1/DDRGK1), and ER translocon machinery (SEC61A1/SEC61G). Importantly, 3’ and 5’ direct capture Perturb-seq produced highly similar relationships (cophenetic correlation with GBC Perturb-seq: *r*=0.95 and *r*=0.95, respectively) and comparable target depletion (median knockdown: 90% for GBC capture, 94% for 3’ sgRNA^cs1^ capture, 95% for 5’ sgRNA capture) to GBC Perturb-seq (**Fig. 2a** and **Supplementary Fig. 3a**). This confirms both the accuracy of direct capture guide identity assignments to cells and the activity of guides with our modified constant regions.

**Figure 2:**
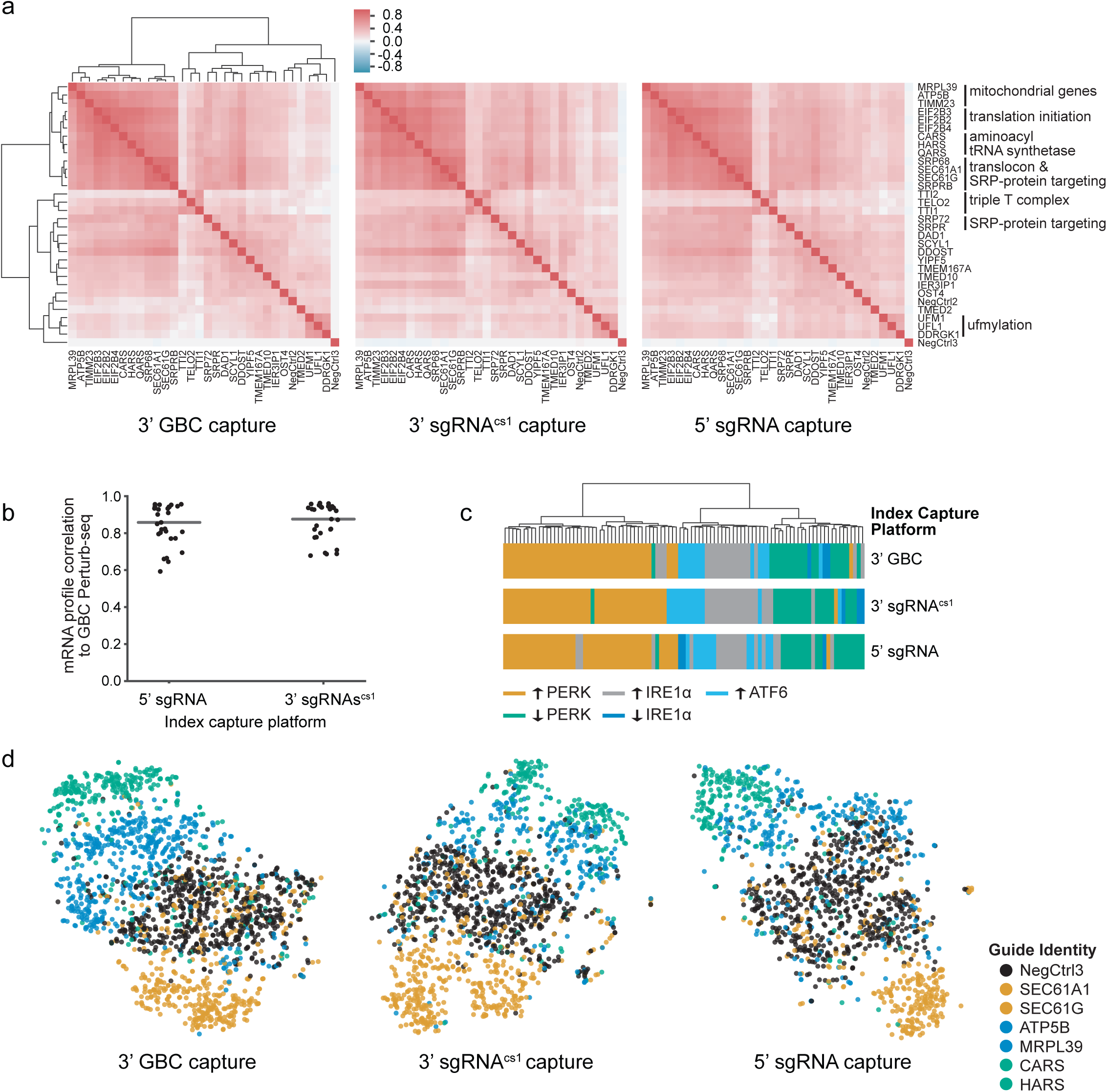
Comparison of direct capture Perturb-seq and GBC Perturb-seq for interrogating gene function and genetic regulation. **(a)** Clustering of perturbations from UPR Perturb-seq experiments conducted with GBC Perturb-seq and direct capture Perturb-seq. Heatmap represents correlations between pseudo-bulk expression profiles for each perturbation. For visual comparison, the rows and columns of all three heatmaps are ordered identically based on the hierarchical clustering of GBC Perturb-seq data. Functional annotations are indicated. **(b)** The correlation of pseudo-bulk expression profiles from direct capture Perturb-seq and GBC Perturb-seq experiments for each perturbation. Profiles were generated from the top 100 differentially expressed genes in GBC Perturb-seq. Grey lines indicate medians. **(c)** Hierarchical clustering of UPR-regulated genes based on co-expression in each of the indicated Perturb-seq experiments. Colors indicate membership in different UPR-regulated groups. **(d)** Single-cell projections based on 10 independent components followed by t-sne. Colors indicate annotation of guide identity.

Next, we assessed the similarity of transcriptional responses produced by each genetic perturbation across platforms. For this, we calculated the expression correlation of the top 100 differentially expressed genes for each perturbation between direct capture and GBC Perturb-seq and found good agreement between platforms (median: *r* = 0.88 for 3’ sgRNA^cs1^ capture, *r*=0.87 for 5’ sgRNA capture) with even higher correlation between perturbations that led to differential expression of >100 genes (**Fig. 2b** and **Supplementary Fig. 2b**). This further supports the accuracy of our guide identity assignments and suggests that our guide-specific RT primers do not globally alter single-cell gene expression profiles. The latter is also supported by the similar number of mRNA molecules and transcripts identified across platforms at a normalized non-saturating sequencing depth (**Supplementary Fig 3c,d**). Lastly, we sought to determine if direct capture Perturb-seq is suitable to elucidate genetic networks. For this, we relied on our prior empirical classification of genes regulated by the three separate signaling branches of the UPR^3^. Examining the covariance of these genes across single cells in our current data, we found gene expression modules that were conserved across platforms (cophenetic correlation: *r*=0.93 for 3’ sgRNA^cs1^ capture, *r*=0.95 for 5’ sgRNA capture), and these modules tended to cluster functionally based on their regulation by the three UPR branches (**Fig. 2c**).

A distinct advantage of Perturb-seq over conventional genetic screens is the ability to reveal and investigate single-cell phenotypes. Thus, we also evaluated the potential of direct capture Perturb-seq to unveil truly single-cell phenomena (e.g. cell-to-cell heterogeneity and bistability). First, we projected cells bearing a subset of the UPR-inducing guides into two dimensions using t-distributed stochastic neighbor embedding (t-sne). In this low-dimensional representation, perturbed cells are distinguishable from unperturbed cells as distinct and functionally-related populations (**Fig. 2d**). Next, we more quantitatively evaluated the single-cell performance of our platforms by training a random forest classifier to classify perturbed and unperturbed (control) cells for each targeted guide. Despite the intrinsic noise of scRNA-seq data, prediction accuracies were highly similar across platforms (correlation with GBC Perturb-seq: r=0.91 for 3’ sgRNA^cs1^ capture, *r*=0.90 for 5’ sgRNA capture) (**Supplementary Figure 3e**).

Cumulatively, our experiments show that direct capture Perturb-seq (on both 3’ and 5’ scRNA-seq platforms) performs comparably to GBC Perturb-seq at the level of both bulk and single cell phenotypes and is therefore well suited for the study of genes and genetic networks. Importantly, these methods are compatible with all common Cas9-derived perturbations, including CRISPRi, CRISPRa, and CRISPR-cutting, and because cs1 and cs2 were not designed for any particular CRISPR system, either approach should be suitable for use with a wide variety of perturbation types. The most promising feature of direct capture Perturb-seq, however, is that it removes the necessity of using indexing transcripts and specialized expression vectors for Perturb-seq, which provides ease of use and adds the flexibility necessary to fulfill the full potential of Perturb-seq.

Importantly, direct capture Perturb-seq will enable single-cell molecular screens with programmed guide combinations. These will have many applications, including efforts to investigate cis-regulatory genome architecture, to elucidate the phenotypes of redundant gene isoforms, and to map genetic interactions in mammalian cells. Notably, traditional growth-based genetic interaction maps are an indispensable tool to objectively and systematically cluster functionally-related genes^18-21^; however, they provide limited molecular insights into the mechanisms underlying interactions. Coupling scRNA-seq with genetic interaction mapping via direct capture Perturb-seq will answer fundamental questions about molecular networks and, more broadly, about the nature of genetic interactions. With this in mind, our initial Perturb-seq implementation (GBC Perturb-seq) was designed to enable the simultaneous transduction of up to three linked guide expression cassettes marked by a single GBC^3^, but as discussed above, interference from lentiviral recombination presents a challenge for scaling GBC Perturb-seq. An alternative way to deliver combinatorial perturbations to cells is super-transduction of single guide expression vectors, such as CROP-seq. However, pairing perturbations through random infection events causes many cells to receive either one or greater than two guides. This results in unwanted sampling of single perturbations and higher-order interactions, as previously discussed^12^. Relying only on capture of active guide molecules, direct capture Perturb-seq presents an obvious solution for scRNA-seq profiling of programmed combinations, one that can be implemented with any vector capable of expressing multiple guides.

Since its inception, Perturb-seq has promised to expand the use of genetic screens in primary cells by reducing the cell engineering and massive cell numbers necessary for pooled screens. Yet these applications have been limited in part by the challenge of integrating traditional Perturb-seq indexing approaches with scRNA-seq techniques used for immunology and developmental biology. Now, direct capture Perturb-seq can be easily multiplexed with the measurement of single-cell V(D)J clonotype, antigen specificity, cell surface protein levels, molecular recording^22^, and transcriptional profiles on the 10x platform with Feature Barcoding technology that was developed in parallel with this work. Additionally, because our 5’ strategy can capture many sgRNA variants including homing sgRNAs, it makes possible the simultaneous measurement of scRNA-seq and developmental lineage in existing mouse strains^23^. We predict that direct guide capture approaches will enable CRISPR-based perturbations to be integrated with other single-cell multi-omic technologies in future work.

Taken together, direct capture Perturb-seq surmounts the limitations of indexing transcript-based platforms to conclusively expand the scale and utility of single-cell molecular screens. We have made our 3’ direct capture Perturb-seq platform accessible through the commercially-available Chromium Single Cell 3’ Reagent Kits v3 with Feature Barcoding technology from 10x Genomics.

## Methods

### Plasmid design and construction

Plasmids for expression of EGFP-NT2 (protospacer: 5’-GACCAGGATGGGCACCACCC-3’)^15^ programmed guide RNAs with capture sequences incorporated into their constant regions were constructed by replacing the constant region sequence in pU6-sgRNA EF1Alpha-puro-T2A-BFP (Addgene, #60955) with modified variants. These variants are versions of the constant region in pU6-sgRNA EF1Alpha-puro-T2A-BFP, originally from sgRNA(F+E)^24^ with a BlpI site^17^, with capture sequences appended to the 3’ end (prior to the terminating poly-T tract) or inserted into the loop region of the so-called “stem loop 2.” Loop modified constant regions also contain an extension to the stem region as found in sgRNA 2.0^16^. Modified sgRNA constant regions in pAX064-pAX099 (**Supplementary Table 1**) were synthesized and inserted using the BstXI and XhoI sites of the parental vector (by Gibson assembly). For pBA896-pBA904 and pBA970 (**Supplementary Table 1**), modified constant regions were synthesized and inserted using BlpI and XhoI. Modified constant regions in pBA893 and pBA894 (**Supplementary Table 1**) was similarly inserted using annealed oligos.

Two vectors evaluated in **Supplementary Figure 1c** (pBA950 labeled as “CROP-seq modified for CRISPRi” and pBA960 labeled as “CROP-seq”) were derived from CROPseq-Guide-Puro (Addgene, #86708). First, an intermediate vector (pBA948) was constructed by replacing the existing selectable marker in CROPseq-Guide-Puro with BFP (synthesized dsDNA inserted by Gibson assembly using PstI and MluI). Then, pBA960 was made by programming the encoded sgRNA with EGFP-NT2, and pBA950 was made by replacing the entire human U6-driven sgRNA expression cassette with the sgRNA expression cassette from pU6-sgRNA EF1Alpha-puro-T2A-BFP (synthesized dsDNA inserted between PpuMI and NcoI by Gibson assembly). This cassette is driven by a modified mouse U6 promoter and encodes a BlpI-containing sgRNA(F+E) programmed with EGFP-NT2. To allow cloning of pBA950, a 21 bp repeat outside of the lentiviral LTRs was also removed.

### Cell culture, DNA transfections, and viral production

RPMI-1640 with 25mM HEPES, 2.0 g/L NaHCO3, 0.3 g/L L-Glutamine supplemented with 10% FBS, 2 mM glutamine, 100 units/mL penicillin and 100 μg/mL streptomycin was used to grow K562 cells. HEK293T cells, used for packaging lentivirus, were grown in Dulbecco’s modified eagle medium (DMEM) in 10% FBS, 100 units/mL penicillin and 100 μg/mL streptomycin. Lentivirus was produced by co-transfecting HEK293T cells with transfer plasmids and standard packaging vectors using *Trans*IT^®^-LTI Transfection Reagent (Mirus, MIR 2306).

### Testing CRISPRi activity of modified guides and guide expression vectors

Guide expression constructs were transduced into GFP+ K562 dCas9-KRAB cells^3^ with centrifugation (2 hours at 1000 x g). Cells were grown without selection for 6-12 days and analyzed by flow cytometry on a LSR-II flow cytometer (BD Biosciences). BFP expression was used to gate for transduced cells. Data in **Supplementary Figure 1c** is reported as the triplicate average of median GFP in cells transduced as indicated normalized to median GFP in cells transduced using pU6-sgRNA EF1Alpha-puro-T2A-BFP. Based on the results of our screen, primers containing our final capture sequences, capture sequence 1 (cs1) (5’-TTGCTAGGACCGGCCTTAAAGC-3’) and capture sequence 2 (cs2) (5’-CCTTAGCCGCTAATAGGTGAGC-3’) were incorporated into gel beads in the Chromium Single Cell 3’ Reagent Kits v3 with Feature Barcoding technology.

### sgRNA library cloning and Perturb-seq screening

We made three CRISPRi libraries, each encoding guide RNAs programmed with 32 unique protospacer sequences (**Supplementary Table 2**) and each with one of three guide constant regions: our parental constant region (as in pU6-sgRNA EF1Alpha-puro-T2A-BFP) or the parental with cs1 inserted into stem loop 2 or cs2 appended to the 3’ end (as described in **Plasmid design and construction**). These libraries target 30 genes that we previously evaluated by Perturb-seq and include two negative controls (NegCtrl2 and NegCtrl3)^3^. We made these libraries by cloning protospacer-containing inserts (annealed oligos from IDT with BstXI/BlpI overhangs) into pBA571, the Perturb-seq GBC library (parental constant region), and two expression vectors, pBA900 (cs2-modified constant region) and pBA904 (cs1-modified constant region) after digestion with BstXI and BlpI. Library vectors were then clonally isolated and verified by Sanger sequencing of the protospacer region and, for pBA571-derived vectors, the corresponding GBC. Notably, pBA571-derived vectors encode guide expression cassettes in the opposite orientation to those in pBA900 and pBA904. Sequence-verified vectors were then individually packaged into lentiviruses, and lentiviral preparations derived from the same parental vector were pooled for co-transduction into cells. To control the representation of guides in cells at the time of scRNA-seq (7 days after transduction), pooling was performed in a manner that would account for both packaging variability and guide effects on cell growth after transduction. This was guided by test infections that determined the percentage of transduced (BFP+) cells remaining after 7 days (quantified by flow cytometry using a LSR-II flow cytometer from BD Biosciences). For each library, pooling was designed to ensure even representation among targeting guides and delivery of NegCtrl2 and NegCtrl3 at 4-fold higher representation.

CRISPRi libraries (constructed with pBA571, pBA900, and pBA904) were spinfected into K562 dCas9-KRAB cells^17^ (2 hours at 1000 x g) with a target multiplicity of infection (MOI) of 0.05. Post centrifugation, cells were transferred to a spinner flask for growth. After 3 days, we measured (by BFP expression) an MOI of 0.04 for the pBA571-derived library, 0.05 for the pBA900-derived library, and 0.05 for the pBA904-derived library, and transduced cells were sorted to near purity on a BD FACSAria2. Seven days post-transduction, cells were separated into droplet emulsions using the Chromium Controller across 5 lanes (10x Genomics). For GBC Perturb-seq, cells transduced with the pBA571-derived library were loaded onto a lane with Chromium Single Cell 3’ Gel Beads v2. For Perturb-seq by 5’ capture of standard sgRNAs, the same cells were loaded onto a lane with Chromium Single Cell 5’ Gel Beads and a 5 pmol spike-in of oJR160 (5’-AAGCAGTGGTATCAACGCAGAGTACCAAGTTGATAACGGACTAGCC-3’) to the RT Master Mix. For Peturb-seq by 5’ sgRNA^cs1^ capture, cells transduced with the pBA904-derived library were loaded onto a lane with Chromium Single Cell 5’ Gel Beads and a 5 pmol spike-in of oJR161 (5’-AAGCAGTGGTATCAACGCAGAGTACTTGCTAGGACCGGCCTTAAAGC-3’) to the RT Master Mix. For Peturb-seq by 3’ sgRNA^cs1^ or sgRNA^cs2^ guide capture, cells transduced with the pBA904- or pBA900-derived libraries, respectively, were loaded onto separate lanes with Chromium Single Cell 3’ Gel Beads v3. For all experiments, cells were loaded to recover ~10,000 cells per lane at a coverage of ~260 cells per guide. Cell viability for all three libraries remained >87% for the duration of the experiment prior to loading onto the Chromium Controller, and cell pools were loaded onto the Chromium Controller at >94% BFP+.

### Sample preparation for sequencing

#### GBC Perturb-seq

The scRNA-seq library for our GBC Perturb-seq screen was prepared according to the Chromium Single Cell 3’ Reagent Kits v2 User Guide (10x Genomics CG00052) with 11 cycles of PCR during cDNA amplification and 11 cycles of Sample Index PCR. Library molecules containing guide barcodes (GBCs) were specifically amplified using KAPA HiFi ReadyMix with 30 ng of the final library as template and 0.6 mM 052-P5 (5’-AATGATACGGCGACCACCGAGATCTACAC-3’) and 0.6 mM of i7 barcoded 055-N708 (5’-CAAGCAGAAGACGGCATACGAGATCCTCTCTGGTCTCGTGGGCTCGGAGATGTGTATAAGAGACAGGACCTCCCTAGCAAACTGGGGCACAAG-3’) (**Supplementary Fig. 1b**). PCR cycling was performed according to the following protocol: (1) 95 C for 3 min, (2) 98 C for 15 s, then 70 C for 10 s (14 cycles) (3) 72 C for 1 min. The resulting GBC sequencing library was purified via a 0.8X SPRI selection.

#### 5’ direct capture Perturb-seq

The 5’ direct capture Perturb-seq libraries were prepared using a protocol modified from the Chromium Single Cell V(D)J Reagent Kits User Guide (10x Genomics CG000086) (**Supplementary Fig. 1b**). The direct capture spike-in oligos oJR160 and oJR161 have an adapter sequence identical to the adapter sequence on the 10x Poly-dT RT Primer (PN-2000007). This servers as a primer binding site for the 10x Non-Poly(dT) primer (PN-220106) during cDNA amplification. Thus, amplification of reverse transcribed guides occurs concurrently with cDNA amplification without an additional spike-in primer. Following 11 cycles of cDNA amplification, library molecules containing guide sequences were enriched by a 0.6X-1.2X double-sided SPRI selection. Separate cDNA fractions were collected from the 0.6X left sided SPRI selection. The cDNA fractions were used to construct scRNA-seq libraries according to the Chromium Single Cell V(D)J Reagent Kits User Guide with 14 cycles of Sample Index PCR. The guide-enriched fractions were quantified by Bioanalyzer High Sensitivity DNA assays (Agilent Technologies, Santa Clara, California) and amplified for sequencing with KAPA HiFi ReadyMix using 5 ng of each library as the template. For 5’ sgRNA capture, guide molecules were amplified using 0.6 mM oJR163 (5’-AATGATACGGCGACCACCGAGATCTACACTCTTTCCCTACACGACGCTCTTCCGATCT-3’) and oJR165 (5’-CAAGCAGAAGACGGCATACGAGATAGGAGTCCGTCTCGTGGGCTCGGAGATGTGTATAAGAGACAGAGTACCAAGTTGATAACGGACTAGCC-3’). For 5’ sgRNA^cs1^ capture, guide molecules were amplified using 0.6 mM oJR163 and oJR166 (5’-CAAGCAGAAGACGGCATACGAGATCATGCCTAGTCTCGTGGGCTCGGAGATGTGTATAAGAGACAGGTACTTGCTAGGACCGGCCTTAAAGC-3’). The PCR cycling protocol consisted of: (1) 95 C for 3 min, (2) 98 C for 15 s, then 70 C for 10 s (12 cycles) (3) 72 C for 1 min. The resulting guide sequencing libraries were purified with a 0.8X SPRI selection. Because the 5’ sgRNA^cs1^ library had a contaminating low-molecular weight species (suspected primer dimers) an additional selection for 248-302 bp fragments was performed using a BluePippin (Sage Science) prior to sequencing.

#### 3’ direct capture Perturb-seq

The 3’ direct capture Perturb-seq libraries were prepared using a protocol modified from the Chromium Single Cell 3’ Reagent Kits v3 User Guide (10x Genomics CG000184) (**Supplementary Fig. 1b**). Following 11 cycles of PCR during cDNA amplification, library molecules containing guide sequences were enriched by a 0.6X-1.2X double-sided SPRI selection. Separate cDNA fractions were collected from the 0.6X left sided SPRI selection. The cDNA fractions were used to construct scRNA-seq libraries according to the Chromium Single Cell 3’ Reagent Kits v3 User Guide with 10 cycles of Sample Index PCR. The guide-enriched fractions were purified by an additional 1X SPRI selection, eluted in 30 uL, and used as the templates for specific amplifications using a nested PCR strategy. For the first PCR step, 5 uL of the guide fractions were mixed with 50 uL Amp Mix (10x Genomics PN#2000047) and 45 uL Feature SI Primers 1 (10x Genomics PN#2000098). The PCR cycling protocol consisted of: (1) 98 C for 45 s, (2) 98 C for 20 s, then 60 C for 5 s, then 72 for 5 s (12 cycles) (3) 72 C for 1 min. The products of these PCRs were then cleaned up using a 1X SPRI selection, eluted in 30 uL, and then used for the second PCR step. Specifically, 5 uL of each library was mixed with 50 uL of Amp Mix (10x Genomics PN#2000047) and 35 uL of Feature SI Primers 2 (10x Genomics PN#2000098) and amplified according to the following protocol: (1) 98 C for 45 s, (2) 98 C for 20 s, then 54 C for 30 s, then 72 for 20 s (5 cycles) (3) 72 C for 1 min. The resulting guide sequencing libraries were cleaned up via a double-sided 0.7X-1.0X SPRI selection.

#### Sequencing

All scRNA-seq and index sequencing libraries were sequenced on 1 lane of a NovaSeq S2. The libraries were sequenced with a custom sequencing strategy: 26 bp of Read 1, 125 bp of Read 2, and 8 bp of Index Read 1. The extended Read 2 was used to sequence protospacers in our 5’ guide sequencing libraries. We note that future users of the Chromium Single Cell 3’ Gel Beads v3 should use a Read 1 length of 28 bp to sequence the entirety of the extended UMIs incorporated into these beads.

#### Data processing and read alignment

We used Cell Ranger 3.0 software (10x Genomics) for alignment of scRNA-seq reads, collapsing reads to unique molecular identifier (UMI) counts, cell calling, and depth normalization of mRNA libraries. Index reads were aligned to expected sequences using bowtie for GBC Perturb-seq and bowtie2 for direct capture Perturb-seq. We observed alignment rates of 0.82 for GBCs, 0.35 for sgRNA^cs2^ protospacers (3’ capture), 0.62 for sgRNA^cs1^ protospacers (3’ capture), 0.71 for protospacers of standard sgRNAs (5’ capture), and 0.62 for sgRNA^cs1^ protospacers (5’ capture). All downstream analyses were performed in Python, using a combination of Numpy, Pandas, scikit-learn, pomegranate, polo, and seaborn libraries.

#### Low cell count / inviable cell removal

Droplet-based scRNA-seq libraries often contain small subpopulations of low UMI count and apoptotic cells. The low UMI count subpopulation likely represents a combination of cells from emulsion droplets that underwent inefficient reverse transcription and droplets that encapsulated fragmented cellular debris. In our experiment with 5’ capture of standard sgRNAs, there was an obvious subpopulation of ~150 (~2%) cells with low mRNA and sgRNA UMI counts. These cells were removed using a threshold. For all experiments, apoptotic cells were removed from single-cell analysis in **Figure 2d** using a random forest regressor that was trained to recognize inviable cells using data from our previously published UPR epistasis experiment^3^.

#### Perturbation identity mapping

During preparation of index read libraries, spurious index-CBC reads can be generated. We attribute these to the encapsulation of ambient GBC-carrying transcripts or guides into emulsion droplets and to PCR chimeras that occur during library preparation for sequencing. To accurately assign guide identities to cells, CBCs marking true guide-expressing cells had to be separated from this background. To compare guide assignments across platforms, we downsampled reads to 25 million aligned indexing reads per GBC or guide sequencing library. At this sequencing depth, saturation of the index libraries is 0.75 for GBC, 0.96 for sgRNA^cs2^ (3’ capture), 0.71 for sgRNA^cs1^ (3’ capture), 0.28 for the library of standard sgRNAs (5’ capture), and 0.60 for sgRNA^cs1^ (5’ capture). For GBC Perturb-seq, we assigned guide identity using a threshold that separates the bimodal distribution of GBC coverage (reads per UMI), as well as UMI and read thresholds, as previously described^3^. For direct capture Perturb-seq, guide coverage distributions were not bimodal at this sequencing depth. Instead, we found that each guide had a bimodal distribution of the number of UMIs per cell (capture rates) and that these capture rates vary across protospacers.

Conceptually, guide capture rates in direct capture Perturb-seq could be influenced by protospacer-dependent variability in guide synthesis and stability, Cas9 binding, and capture or template switching efficiency. To robustly assign guide identities with varying protospacer capture rates, we fit a mixture model to the distribution of UMIs per cell for each protospacer to separate cells expressing the guide of interest (upper mode) from background cells (lower mode). Specifically, for each protospacer, we used expectation maximization to fit a two-component mixture model consisting of a Poisson (background) and Gaussian (guide-expressing) distribution to the log_2_ transformed UMIs per cell (representative model in **Figure 1g**). These mixture models allow for probabilistic assignment of cells to background populations or guide-expressing populations, where each cell with a posterior probability >0.5 of belonging to the upper mode component is assigned a guide identity. This procedure produced a coherent proportion of cells assigned to each guide identity in the libraries (**Supplementary Figure 2b**) and a coherent multiplet rate across platforms (**Fig. 1i**). Unless otherwise indicated, only cells with a single assigned guide were considered for downstream analysis. Guide capture rates were significantly correlated across direct capture platforms (*p*<0.001 for each pairwise comparisons of platforms) (**Supplementary Fig. 2c**). Notably, we observed a significant relationship between guide capture rate (UMIs per cell) and the penultimate 5’ terminal nucleotide of the protospacer (all protospacers begin with a guanine that is necessary for transcription at the U6 promoter) (Kruskal-Wallis H-test: *p*<0.05 for each platform) with the highest capture rates for guanine (**Supplementary Fig. 2d,e**). We did not observe a significant relationship between total protospacer GC content and the guide capture rate (**Supplementary Fig. 2f**).

#### Expression normalization, average expression profiles, and target knockdown

We normalized for differences in mRNA capture and coverage across cells by rescaling each cell to have the same median total UMIs (i.e., each row of the raw expression matrix is rescaled to have the same sum). Then, expression of each gene was z-normalized with respect to the mean and standard deviation of that gene in the control cell population bearing NegCtrl3. For **Figures 2a and 2b**, we generated pseudo-bulk RNA-seq profiles by averaging the normalized expression profiles of well-expressed genes (excluding genes with a mean expression <1 UMI per cell for **Figure 2a** and <0.5 UMI per cell for **Figure 2b**) for all cells expressing a guide, excluding cells assigned multiple guide identities.

#### Hierarchical clustering of genes and cophenetic correlation

For each library, we used the pseudo-bulk RNA-seq profiles to calculate a guide-guide Spearman’s rank correlation matrix, to which we applied Ward’s method to hierarchically cluster genes. We optimally ordered hierarchical clusters to minimize the distance between successive leaves. For **Figure 2a**, all heatmaps are ordered based on the GBC Perturb-seq clustering to enable visual comparison of the correlation matrices. To quantitatively compare the similarity of the correlation matrices, we calculated a cophenetic correlation, the correlation of all pairwise similarities between perturbation profiles across platforms.

#### Correlation of average expression profiles

For **Figure 2b**, we used random forest classifiers to identify differentially expressed genes. Specifically, for each guide, we trained a random forest classifier (scikit-learn extremely randomized trees with 1000 trees in the forest) to predict perturbation status, using the normalized expression profile of each cell for genes with a mean expression >0.5 UMI per cell as features. For each guide, the top 100 genes whose expression level could be used to separate perturbed and unperturbed cells in GBC Perturb-seq were considered differentially expressed. Using these sets of differentially expressed genes, we calculated the correlation of pseudo-bulk RNA-seq profiles for each perturbation between direct capture Perturb-seq and GBC Perturb-seq (**Figure 2b**).

The advantage of the above approach is that we assess the similarity of average expression profiles across platforms regardless of the strength of the perturbation because we do not employ a strict cutoff. However, as this strategy always returns 100 genes per perturbation without regard for statistical significance, it may return genes that are not truly differentially expressed for those perturbations that do not have a robust transcriptional phenotype, which may deflate the apparent average correlation between platforms. In order to account for this, for each perturbation we performed a two-sample Kolmogorov–Smirnov test for each gene and considered genes differentially expressed when the Benjamini/Yekutieli FDR-corrected *p*<0.05 for GBC Perturb-seq. As expected, we found that the number of differentially expressed genes for each perturbation was associated with the degree of correlation between the pseudo-bulk expression profiles for that perturbation between direct capture Perturb-seq and GBC Perturb-seq (Spearman’s rank correlation: ρ=0.84, *p*=2e-9 for 3’ sgRNA^cs1^ capture; ρ=0.86, *p*=5e-10 for 5’ sgRNA capture) (**Supplementary Figure 3b**).

#### Comparison of clustering of UPR-regulated genetic network

Previously, we used Perturb-seq to dissect UPR-mediated transcription into sets of genes regulated by the UPR signaling branches controlled by IRE1α, PERK, and ATF6^3^. We expected these gene sets to be differentially expressed by our UPR-inducing guide RNAs and to meaningfully covary based on differential UPR branch activation. Therefore, we reasoned that this gene set would serve as a useful test case for the ability of direct capture Perturb-seq to cluster genetic regulons. To this end, we used the normalized expression profiles of all cells from our GBC, 3’ sgRNA^cs1^, and 5’ sgRNA Perturb-seq experiments to calculate gene-gene Spearman’s rank correlation matrices for this gene set. We quantitatively assessed the similarity of the clustering produced by calculating the cophenetic correlation, the correlation of all pairwise similarities between genes across platforms. As a visual aid, we applied Ward’s method to the coexpression matrix to hierarchically cluster genes, where the dendrograms tend to place coregulated genes near one another. The genes were roughly grouped based on their branch activation, in agreement with our previous work.

#### Single cell analysis

To determine the single cell performance of direct capture Perturb-seq, we again employed random forest classifiers. For strong genetic perturbations, we generally expected perturbed cells to be transcriptionally distinguishable from unperturbed cells. For each guide, we split our data into training (80%) and testing (20%) data and trained a random forest classifier on normalized expression profiles to separate perturbed and unperturbed (NegCtrl3) cells. We tested the accuracy (balanced for perturbed and unperturbed cells) of our classifiers on the remaining 20% of cells. In this regime, the accuracies of our random forest classifiers can serve as proxies for the single-cell performance of Perturb-seq across platforms. We used a Wilcoxon signed-rank test to test for differences in classification accuracy across platforms and failed to reject the null hypothesis that direct capture Perturb-seq performs comparably to GBC Perturb-seq (Wilcoxon singed-rank test: *p*=0.2 for 3’ sgRNA^cs1^ capture; *p*=0.6 for 5’ sgRNA capture) (**Supplementary Figure 3e**).

To visually inspect single cell performance, we took a subset of cells bearing guides producing robust perturbations, specifically those targeting SEC61A1, SEC61G, ATP5B, MRPL39, CARS, HARS, and NegCtrl3. We reduced dimensionality by computing 10 independent components (using FastICA) followed by t-distributed stochastic neighbor embedding (t-sne) to project cells into two dimensions.

## Acknowledgements

We thank members of the Weissman lab, 10x Genomics, Eric Chow, and Rene Sit for helpful discussions. This work was funded by National Institutes of Health Grants P50 GM102706, U01 CA168370, R01 DA036858 (all to J.S.W.) as well as T32 GM007618 (J.M.R.) and the Chan Zuckerberg Initiative. J.S.W. is a Howard Hughes Medical Institute Investigator. T.M.N. is a fellow of the Damon Runyon Cancer Research Foundation (T.M.N DRG-[2211-15]).

## Author Contributions

J.M.R., T.M.N., J.S.W., and B.A. conceived, designed, and interpreted the experiments and wrote the manuscript. J.M.R., A.X., and B.A. designed, built, and validated modified guide constant regions, expression vectors, and libraries. J.M.R. performed and A.X. and B.A. contributed to Perturb-seq experiments. J.M.R. performed data analysis with support from T.M.N. D.R. designed the library of candidate capture sequences. 10x Genomics with E.J.M, J.M.T, D.R., N.S., and T.S.M designed and built the Chromium Single Cell 3’ Reagent Kits v3 with Feature Barcoding technology.

## Supplementary Information

**Supplementary Table 1: Capture sequences and sgRNA expression vectors**.

**Supplementary Table 2: sgRNA and GBC library sequences**.

**Supplementary Figure 1:**
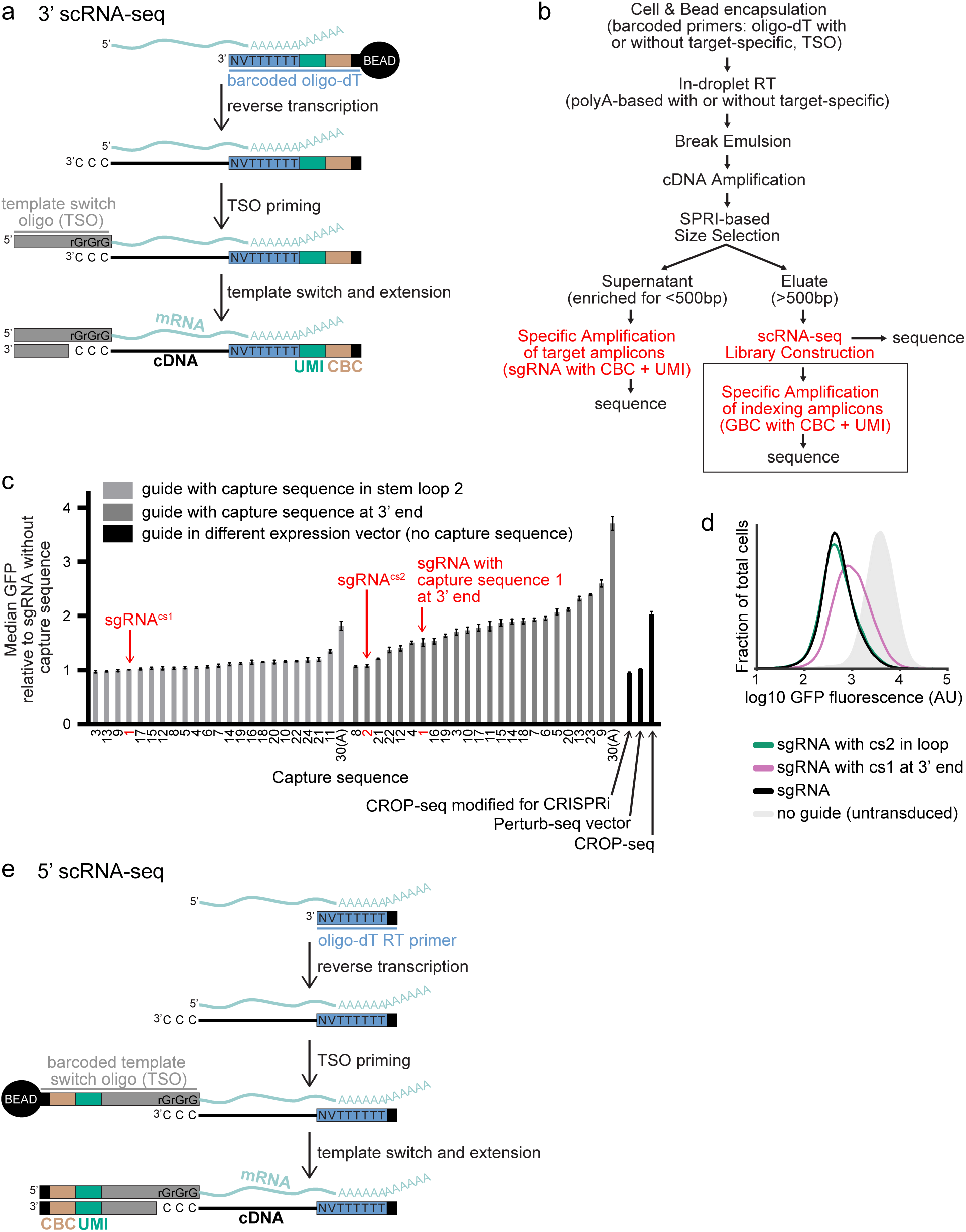
Optimized modification of guide constant regions to enable 3’ direct capture Perturb-seq. **(a)** Schematic of 3’ single-cell RNA-sequencing (3’ scRNA-seq). Polyadenylated mRNAs from individual cells (top, light blue) anneal to barcoded oligo-dT primers in emulsion droplets (delivered to droplets on gel beads) and are reverse transcribed into indexed cDNA (bottom). TSO, template switch oligo. UMI, unique molecular identifier. CBC, cell barcode. **(b)** Schematic of experimental workflows for GBC or direct capture Perturb-seq (3’ or 5’) based on protocols from 10x Genomics. Red indicates generation of sequencing libraries. Box details construction of index sequencing library for GBC Perturb-seq, which is based on our previously published protocol^3^. **(c)** CRISPRi activity of guides carrying the indicated capture sequences (all programmed with the GFP-targeting protospacer EGFP-NT2) in GFP+ K562 dCas9-KRAB cells 6 or 10 days post-transduction. Data from guides selected for direct capture experiments (sgRNA^cs1^ and sgRNA^cs2^) are indicated. 30**(A)** indicates 30 adenines. For comparison, data from standard guides also programmed with EGFP-NT2, without capture sequences, and expressed from 3 other vectors (indicated) were also included. One of these, indicated as “CROP-seq,” has a different (previously published)^4^ constant region and is expressed from a different promoter. Data was collected in three independently controlled experiments and represents the average of triplicate measurements normalized to control measurements ± standard error. For reference, the median GFP of transfected controls (relative to the median of untransfected cells in the same well) were 0.14, 0.14, and 0.11. **(d)** Gaussian kernel density estimates of normalized flow-cytometry measurements representing GFP expression demonstrate CRISPRi activity of the indicated guide RNAs (programmed with the GFP-targeting protospacer EGFP-NT2). The negative and positive control data are the same as in **Figure 1d**. **(e)** Schematic of 5’ single-cell RNA-sequencing (5’ scRNA-seq). Polyadenylated mRNAs from individual cells (top, light blue) anneal to unbarcoded oligo-dT primers in emulsion droplets (delivered to droplets as free oligos) and are reverse transcribed. Indexing of cDNA (bottom) occurs after template switching allows for extension of barcoded TSOs (delivered to droplets on gel beads).

**Supplementary Figure 2:**
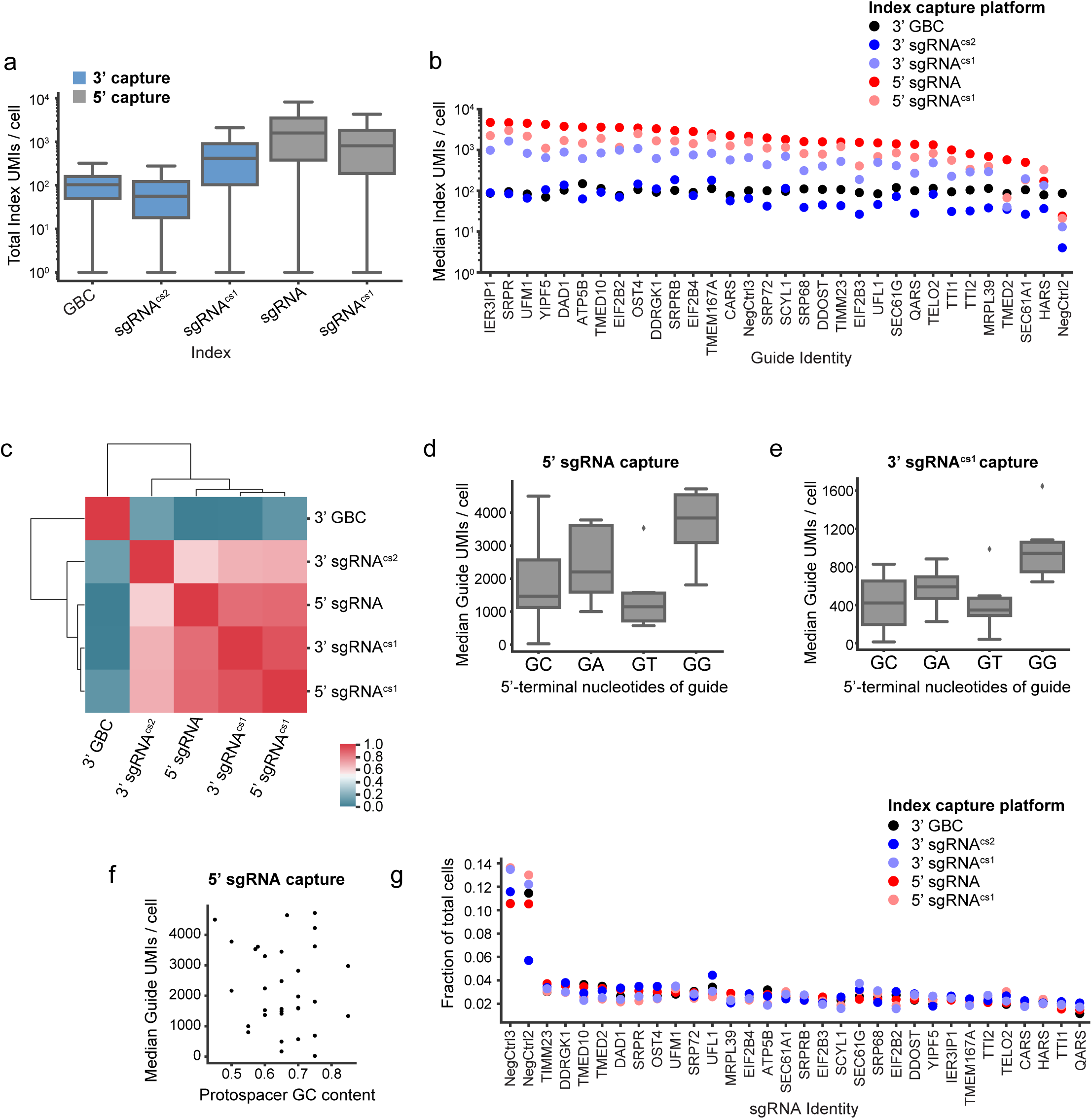
Cell indexing by direct guide capture is robust and comparable to indexing by GBC capture. **(a)** Boxplot of total index (GBC or guide) UMI counts per cell for all cells (prior to guide identity mapping). Upper and lower whiskers display the highest and lowest data points within 1.5 interquartile range of the third and first quartiles, respectively. **(b)** Median index UMI counts per cell (capture rate) for cells assigned to each guide identity in Perturb-seq experiments (n=32 guides per experiment). Across platforms, NegCtrl2 has the worst capture rate, which may be explained by the fact that this negative control guide has a protospacer containing an extended run of guanine nucleotides (5’-GCGATGGGGGGGTGGGTAGC-3’). Data plotted here are also plotted in **Figure 1h**. **(c)** For each pairwise comparison of Perturb-seq experiments, we calculated a correlation of guide capture rates. Across experiments performed with direct capture Perturb-seq, guide capture rate is highly correlated (*r*>0.6), suggesting that protospacer-dependent features influence guide capture. **(d)** Boxplot of median guide UMIs per cell stratified by the 5’ terminal nucleotides of the protospacer. The displayed data is from Perturb-seq by 5’ sgRNA capture; for all platforms, there is a significant relationship between capture rate and the 5’ terminal nucleotides (Kruskal-Wallis H-test: *p*<0.05 for each platform). **(e)** Boxplot of median guide UMIs per cell stratified by the 5’ terminal nucleotides of the protospacer. The displayed data is from Perturb-seq by 3’ sgRNA^cs1^ capture; for all platforms, there is a significant relationship between capture rate and the 5’ terminal nucleotides (Kruskal-Wallis H-test: *p*<0.05 for each platform). **(f)** Scatterplot of the median number of UMIs per cell and protospacer GC content for each of 32 guides. The displayed data is from Perturb-seq by 5’ sgRNA capture; for each platform, we observed no significant correlation between capture rate and protospacer GC content (*p*>0.4 for each platform). **(g)** Identity assignment rates per guide for GBC Perturb-seq and direct capture Perturb-seq experiments. Balanced representation among cells assigned to each of 32 guides (with intentionally 4-fold overrepresented negative controls) was achieved by titering lentiviruses prior to pooling (see Methods).

**Supplementary Figure 3:**
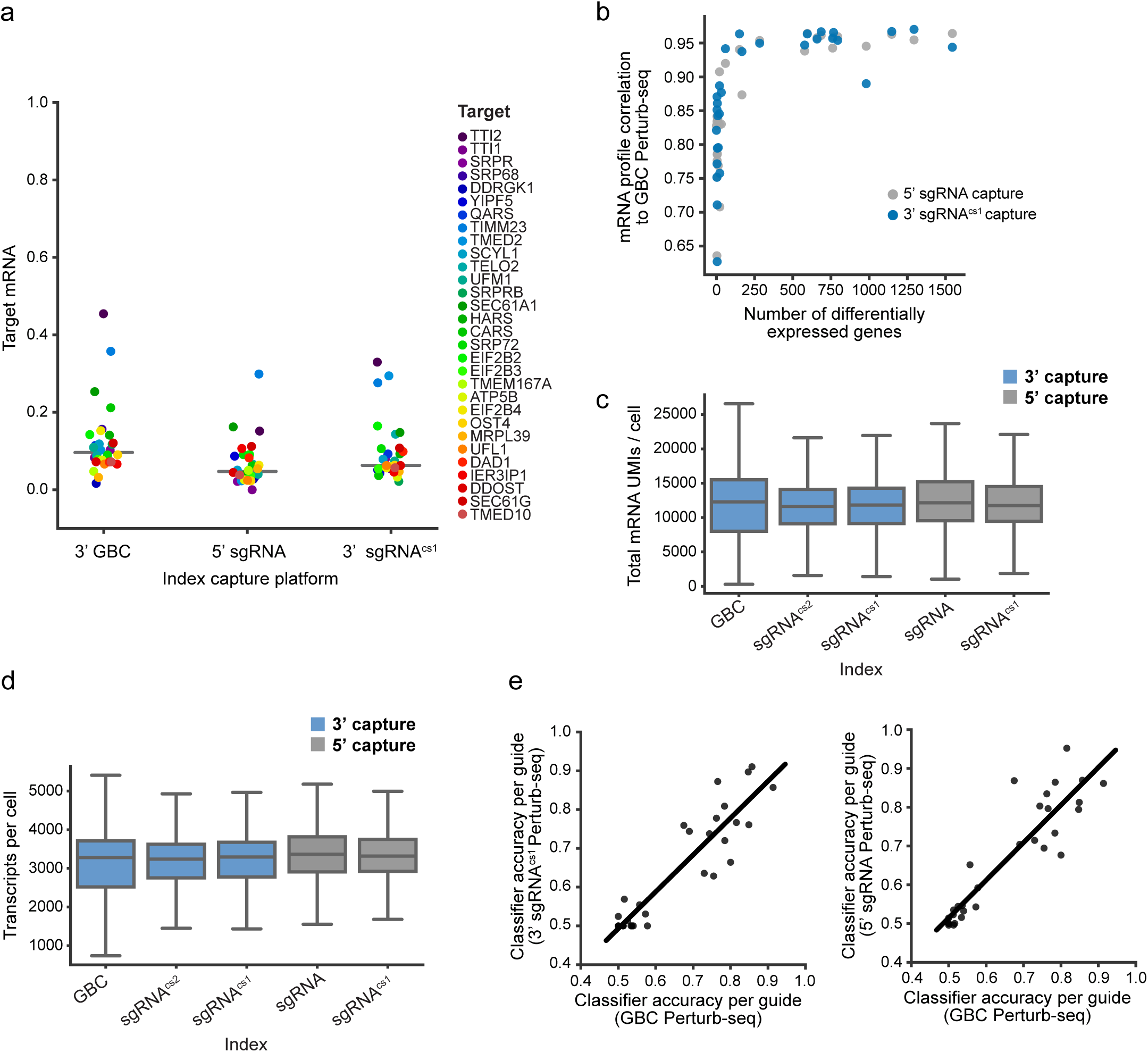
Direct capture Perturb-seq performs comparably to GBC Perturb-seq for phenotypic analysis. **(a)** Mean target knockdown (fraction mRNA remaining plotted) for each targeting guide (n=30) in the indicated experiments. For each guide, the data point represents the mean normalized expression level of the target gene across cells bearing the corresponding guide divided by the mean normalized expression level of the target gene in control cells (NegCtrl3). **(b)** Scatterplot of the relationship between the number of differentially expressed genes for each guide (determined by a KS test using GBC Perturb-seq data) and the correlation of pseudobulk expression profiles between GBC Perturb-seq and direct capture Perturb-seq on the indicated platform. **(c)** Boxplot of the number of mRNA UMIs detected per cell for each Perturb-seq experiment after sequencing depth normalization such that all libraries have the same number of reads per cell. **(d)** Boxplot of the number of mRNA transcripts detected per cell for each Perturb-seq experiment after sequencing depth normalization such that all libraries have the same number of reads per cell. **(e)** Scatterplots of the balanced accuracy of random forest classifiers trained to distinguish perturbed and unperturbed (NegCtrl3) cells for each guide on the indicated platforms. Direct capture Perturb-seq accuracies were highly correlated with GBC Perturb-seq (correlation: *r*=0.91 for 3’ sgRNA^cs1^ capture; *r*=0.90 for 5’ sgRNA capture). We failed to detect significant differences in performance between direct capture Perturb-seq and GBC Perturb-seq (Wilcoxon signed-rank test: *p*=0.2 for 3’ sgRNA^cs1^; *p*=0.6 for 5’ sgRNA capture).

